# Osteoarthritic chondrocytes exert higher contractile forces and exhibit enhanced protrusive activity when cultured in 3D degradable hydrogels

**DOI:** 10.1101/2025.02.27.640613

**Authors:** Maxim Vovchenko, Nele Vaes, Jorge Barrasa-Fano, Apeksha Shapeti, Rocío Castro-Viñuelas, Laurens Kimps, Christ Glorieux, Ilse Jonkers, Hans Van Oosterwyck

**Author notes:** equal contribution (Maxim Vovchenko, Nele Vaes).

## Abstract

Osteoarthritis (OA) induces phenotypic changes in chondrocytes as well as alterations in matrix composition and mechanics. Yet, its impact on active cell-generated forces, a key indicator of cell-matrix interaction, remains poorly characterized. In this study, we systematically compared the force generation capacity and associated proteins of interest between human OA and non-OA articular chondrocytes and how they are affected by cell culture dimensionality (2D versus 3D) and matrix degradability. Using traction force microscopy (TFM) combined with high-resolution immunostainings we show that OA alters the expression and organization of proteins involved in force exertion and transmission across both 2D and 3D cultures, but only in a 3D degradable hydrogel environment do these changes translate into higher cell-generated contractile forces This increased force generation correlates with elevated protrusive activity, higher actomyosin content and engagement, as well as altered localization of adhesion and matrix proteins, all of which could contribute to increased cell-matrix interaction in OA chondrocytes. In contrast, OA chondrocytes display no increase in cell tractions when cultured on 2D hydrogel substrates. These findings demonstrate that the detection and interpretation of OA-related alterations in chondrocyte mechanobiology are strongly dependent on the dimensionality and degradable properties of the culture system. Our results highlight the critical role of 3D degradable environments in revealing disease-associated changes in chondrocyte force generation and emphasize the necessity of carefully selecting model systems when investigating OA mechanobiology.

## Introduction

Osteoarthritis (OA) is a degenerative joint disease characterized by progressive and irreversible articular cartilage breakdown, subchondral bone remodeling, and synovial inflammation, leading to pain and loss of joint function. OA is associated with a hypertrophic chondrocyte phenotype marked by a disruption in the balance between anabolic matrix synthesis and catabolic degradation [1]. This includes increased expression of matrix metalloproteinases (MMPs) [2], [3], [4] and the secretion of pro-inflammatory cytokines such as tumor necrosis factor (TNF)-α, interleukin (IL)-1α, IL-1β, IL-6 and IL-8, which further stimulate the production of catabolic enzymes [5].

Mechanical cues are known to play a critical role in cartilage homeostasis and OA pathogenesis. In healthy cartilage, moderate, cyclic mechanical loading stimulates the synthesis of extracellular matrix components and promotes interstitial fluid flow for nutrient transport [6]. Conversely, both excessive and insufficient joint loading can disrupt healthy chondrocyte function, leading to matrix degradation, elevated MMP levels and increased inflammation [7], [8], [9], [10]. Chondrocytes sense and respond to mechanical stimuli through a process known as mechanotransduction, which involves various mechanosensory molecules and complexes, including integrins, the actin cytoskeleton, ion channels, and primary cilia. In OA, this mechanosensing system becomes dysregulated, exacerbating disease progression in response to mechanical loading.

Integrins and the actin cytoskeleton are particularly important mechanosensors for transducing mechanical signals between the extracellular (pericellular) matrix and the intracellular environment. In addition to mediating cell adhesion and force transmission, OA-associated changes in integrin signaling such as dysregulated αVβ3 signaling have been linked to increased inflammation and cartilage degradation [11]. OA also impacts actin organization in chondrocytes with an observed decrease in cortical F-actin [12]. Beta1 integrins are among the most relevant integrin subtypes in chondrocytes. However, their expression in OA remains incompletely understood, with conflicting findings across species and models. Some rodent studies report either an increase or decrease in β1 integrin levels, while primate models suggest an increase, and human explants show variable results. For example, two studies found no significant change in β1 integrin expression in chondrocytes within human OA explants compared to healthy explants [13], [14], while another reported that a decrease in β1 integrin in isolated OA chondrocytes correlated with increasing cartilage damage severity [15].

Given the effect of OA on the actin cytoskeleton and integrins in chondrocytes, we hypothesized that OA may influence a chondrocyte’s ability to actively generate forces through dysregulated actomyosin contractility and/or cell—extracellular matrix (ECM) interactions. Interestingly, increased phosphorylated myosin light chain (pMLC) levels observed in both human and mouse OA cartilage could suggest aberrant force generation in disease states [16]. While active, actomyosin-driven cellular forces are known to play a critical role in mechanotransduction, their contribution in OA remains unexplored. These intrinsic forces not only make use of the same molecular machinery for force sensing and transmission as external forces do, but they can also modulate how external mechanical signals are transmitted from the extracellular to the intracellular environment [17], [18]. Thus, OA-related changes in active force generation may contribute to dysfunctional mechanotransduction in response to external mechanical loading, thereby leading to disease progression.

Previous studies have used 2D traction force microscopy (TFM) to investigate how factors such as substrate stiffness, soluble cues [19], external loading, vimentin expression [20], and shear stress [21] influence active force generation by healthy chondrocytes. However, these studies were conducted in 2D culture systems on soft hydrogels, which do not capture the native 3D cartilage environment. While chondrocyte phenotype is better preserved in 3D systems [22], [23], [24], culture dimensionality likely also affects cytoskeletal organization, integrin-mediated cell-ECM interactions, and active force exertion. Moreover, these studies focused exclusively on healthy cells and did not examine OA-related changes in chondrocyte active force exertion. Therefore, extending active force measurements to OA chondrocytes in 3D hydrogels using 3D TFM is both necessary and more physiologically relevant.

Given these knowledge gaps, we investigated the impact of OA on chondrocyte force generation using TFM on cells isolated from healthy and OA human cartilage. To assess the influence of culture dimensionality, we performed both 2D TFM for chondrocytes on non-degradable polyacrylamide (PAA) gels and 3D TFM for chondrocytes embedded in degradable polyethylene glycol (PEG) hydrogels. This dual approach allowed us to evaluate how the cell microenvironment impacts OA-related mechanical and mechanobiological changes. Force measurements were complemented with confocal microscopy imaging to assess molecular and structural features underlying force generation. Specifically, we examined cell and nuclear morphology, along with key cytoskeletal (actin, pMLC), adhesional (β1 integrin), and matrix (fibronectin) proteins to link changes in forces to OA-associated molecular alterations.

## Results

### 1. OA chondrocytes are larger, exhibit altered actin and integrin organization and deposit more matrix when cultured on 2D substrates

We cultured primary human articular chondrocytes isolated from hip joints of non-OA and OA donors (3 patients for each condition) on soft (Young’s modulus 4.5 kPa) 2D fibronectin-coated non-degradable PAA gels. We chose this stiffness to avoid inducing a catabolic shift in non-OA cells as previously reported for stiff matrices [16]. Sparse cell seeding, mirroring the natural sparse *in vivo* distribution of chondrocytes, allowed us to assess individual cell behavior—more specifically, changes in cell morphology, components of cytoskeletal organization and cell-matrix adhesion, namely F-actin, β1 integrin and fibronectin, that are known to be altered upon OA. To determine if this phenotypic shift persist when OA chondrocytes are cultured on soft matrices, we performed immunostaining for these molecules after 24 hours of culture.

OA chondrocytes exhibited a significantly larger projected cell and nucleus area compared to non-OA cells (Fig. 1a, 1b), consistent with hypertrophic changes associated with OA progression [4] [25]. No difference in cell morphology metrics (solidity, eccentricity and circularity) was observed between non-OA and OA chondrocytes (Fig. S1a and Fig. S4). Moreover, OA chondrocytes did not show a difference in protrusive activity compared to non-OA cells (Fig. S1b and Fig. S6). The larger area was not accompanied by an increase in total F-actin (Fig. 1d, left) or β1 integrin expression (Fig. 1c and 1f, left). Instead, the combination of unchanged protein expression and increased cell area resulted in a general decrease in the density of both actin (Fig. 1d, middle) and β1 integrin (Fig. 1f, middle). While the expression of these proteins is not affected by OA, we observed moderate differences in distribution. Previously a reduction in cortical actin in OA cells has been reported [26]. Although not significant, OA cells demonstrated a trend for lower cortical levels than non-OA cells (Fig. 1d, right). OA cells also redistributed their β1 integrin content, exhibiting significantly reduced localization at the cell cortex (Fig. 1f, right). OA cells deposited more fibronectin than non-OA cells (Fig. 1e and 1g, left). This increased deposition was proportional to the change in size, since the amount of fibronectin normalized to cell area remained unchanged (Fig. 1g, middle). Additionally, we observed that OA cells had less preference for depositing fibronectin at the cortical region than non-OA cells (Fig. 1g, right). Interestingly, deposited fibronectin was sometimes organized into distinct strand-like patterns (Fig. 1e, top right), similar to the observations from a previous study [27]. This organization was observed in 56% of OA chondrocytes compared to only 16% of non-OA cells.

**Figure 1.**
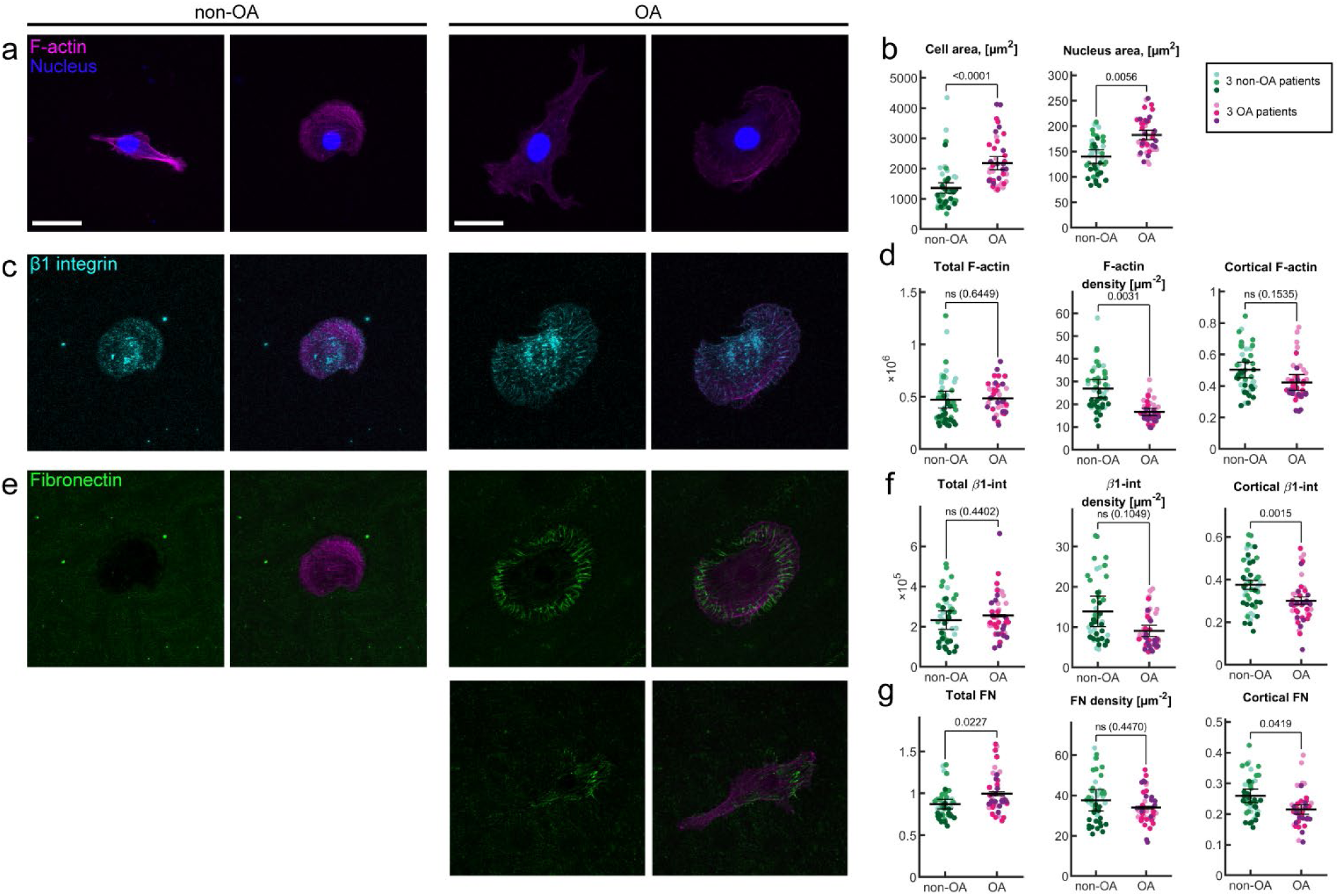
2D immunostaining analysis of cytoskeletal organization and cell-matrix adhesion in non-OA and OA chondrocytes. **a** Maximum intensity projections of F-actin expression (magenta) and nucleus (blue) of two representative non-OA (healthy) and OA chondrocytes, 24h after seeding on a polyacrylamide (PAA) hydrogel. **b** Cell and nucleus area. **c** Maximum projection of β1 integrin expression (cyan) and overlay with F-actin (magenta) for a representative non-OA and OA chondrocyte. **d** Quantification of F-actin expression: Total F-actin intensity of a cell, F-actin density (computed as Total F-actin intensity normalized by cell area), Cortical F-actin (computed as ratio of F-actin present in the cortex normalized by the total F-actin intensity). **e** Fibronectin expression (green) and overlay with F-actin (magenta) for a representative non-OA chondrocyte and two representative OA chondrocytes with different fibronectin expression patterns. **f** Quantification of β1 integrin expression: Total β1 integrin intensity of a cell, β1 density (computed as Total β1 integrin intensity normalized by cell area), Cortical β1 integrin (computed as ratio of β1 integrin present in the cortex normalized by the total β1 integrin intensity). **g** Quantification of Fibronectin expression: Total Fibronectin (computed as Total Fibronectin intensity normalized to background signal), Fibronectin density (computed as Total Fibronectin intensity normalized by cell area), Cortical Fibronectin (computed as the ratio of fibronectin present in the cortex normalized by the Total Fibronectin intensity). All statistical comparisons were performed using generalized linear mixed-effects models to account for the nested structure of the data. Error bars represent SEM, scalebars are 30 µm. Each green and magenta hue in the plots corresponds to a different patient (N=3 for each condition), each dot represents a single cell measurement (n≥15 for every patient).

### 2. OA chondrocytes in 2D show no difference in force generation capacity compared to non-OA chondrocytes despite alterations in molecular organization

We then performed 2D TFM experiments for the same PAA substrates as described in the previous section with fluorescent nanobeads (200 nm diameter) incorporated in the gels to allow for 2D TFM 24 hours after cell seeding (Fig. 2a). We quantified cell-induced PAA displacements, from which cell-generated tractions were inferred based on Tikhonov regularized Fourier transform traction cytometry. We did not observe any differences in the magnitude of maximum PAA displacements, tractions or global contractility between OA and non-OA chondrocytes (Fig. 2b, 2c and 2d, left). Similarly, traction distribution was not affected either, with most of the high tractions being exerted at the cortical area for both OA and non-OA chondrocytes (Fig. 2d, right).

**Figure 2.**
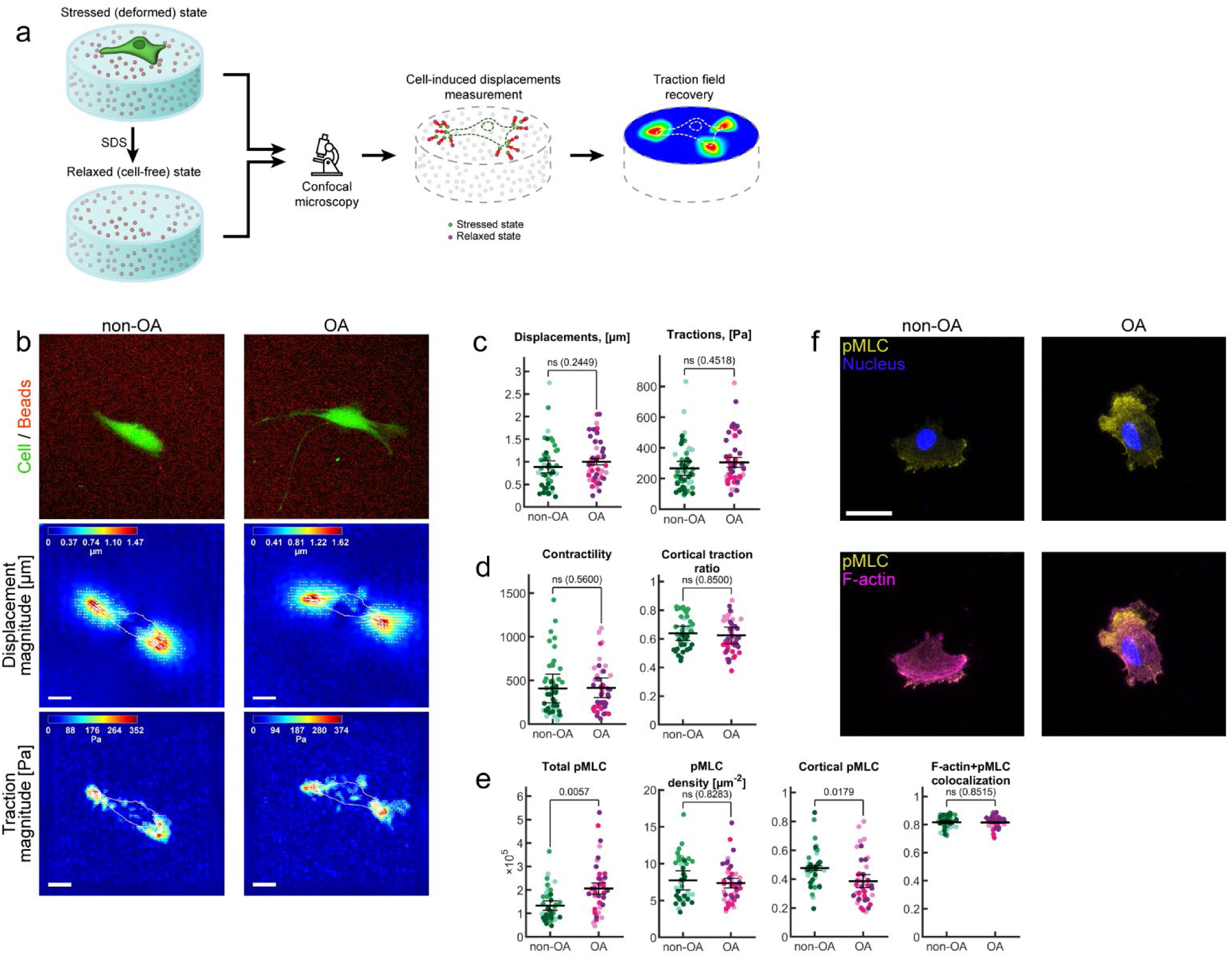
2D TFM analysis of cell tractions and contractility in OA and non-OA chondrocytes. **a** 2D TFM workflow. **b** 2D TFM results illustration. (top) Representative maximum projected images of beads (red) and chondrocyte (green) for one non-OA and one OA chondrocyte. (middle) Corresponding displacement magnitude. (bottom) Corresponding traction magnitude. **c** 99^th^ percentile bead displacement magnitude [µm] and the 99^th^ percentile traction magnitude [Pa]. **d** Contractility, defined as the sum of the (signed) traction vector components (positive if tensile, negative if compressive) along the axis that connects the corresponding pixel to the cell centroid, Quantification of cortical traction ratio as the ratio between tractions in the cortical area and total tractions. **e** Total pMLC intensity of a cell, pMLC density (computed as total pMLC intensity normalized by the cell area), Cortical pMLC (computed as ratio of pMLC intensity in the cell cortex normalized by total pMLC intensity), F-actin and pMLC colocalization. **f** Maximum intensity projection of pMLC expression (yellow) and overlay with F-actin (magenta) for a representative non-OA and OA chondrocyte. All statistical comparisons were performed using generalized linear mixed-effects models to account for the nested structure of the data. Error bars represent SEM, scalebars are 30 µm. Each green and magenta hue in the plots corresponds to a different patient (N=3 for each condition), each dot represents a single cell measurement (n≥15 for every patient).

Immunostaining revealed elevated total phosphorylated myosin light chain (pMLC) levels in OA chondrocytes (Fig. 2e, left, and 2f), in line with *in vivo* observations on OA cartilage tissue [16]. This increase in pMLC was proportional to the observed increase in cell area for OA chondrocytes, resulting in similar pMLC densities for OA and non-OA cells (Fig. 2e, second panel). Similar to the distribution of F-actin, β1 integrin and fibronectin, cortical pMLC levels in OA cells were reduced (Fig. 2e, third panel). Actin-pMLC colocalization was comparable between OA and non-OA chondrocytes (Fig. 2e, right), suggesting that actomyosin coupling remained intact. Together with the findings of the previous section, it means that OA does not affect traction generation by chondrocytes when cultured on 2D substrates, despite the observed differences in terms of total pMLC (increased for OA chondrocytes) and distribution of actin, pMLC, β1 integrins and fibronectin (less cortical for OA chondrocytes).

### 3. OA chondrocytes in 3D exhibit a more protrusive phenotype than non-OA chondrocytes

To investigate chondrocyte behavior in a more physiologically relevant environment, we cultured cells in 3D soft (Young’s modulus 520 Pa) degradable PEG hydrogels with MMP-cleavable peptides (sensitive to MMP-1 and MMP-3, which are both upregulated upon OA). Functionalization of the hydrogels with RGD peptides (as found in fibronectin) facilitated adhesion through RGD-binding integrins, including β1-containing subtypes such as α5β1, which are broadly expressed in chondrocytes. Consistent with our findings in 2D culture, OA chondrocytes exhibited a significantly larger projected cell area (indicative of increased cell volume) (Fig. 3b, left), but unchanged nuclear size (Fig. 3b, right). Other global cell morphology metrics (cell solidity, cell eccentricity and cell circularity) were not affected by OA (Fig. S2). Interestingly, OA cells displayed a more protrusive phenotype than non-OA cells, as characterized by a higher number of protrusions, increased protrusion length and area (Fig. 3d, 3e). To determine whether OA-related morphological changes are coupled to cytoskeletal changes, we further quantified F-actin intensity levels and localization. In contrast to our 2D findings, OA chondrocytes exhibited significantly higher total F-actin levels (Fig. 3c, left). This increase was not proportional to the increase in cell size, since F-actin density was also significantly higher compared to non-OA chondrocytes (Fig. 3c, middle). Interestingly, F-actin distribution analysis showed that for OA chondrocytes F-actin is more localized in protrusions than for non-OA cells, based on a higher protrusion-to-cell body F-actin intensity ratio (Fig. 3c, right).

**Figure 3.**
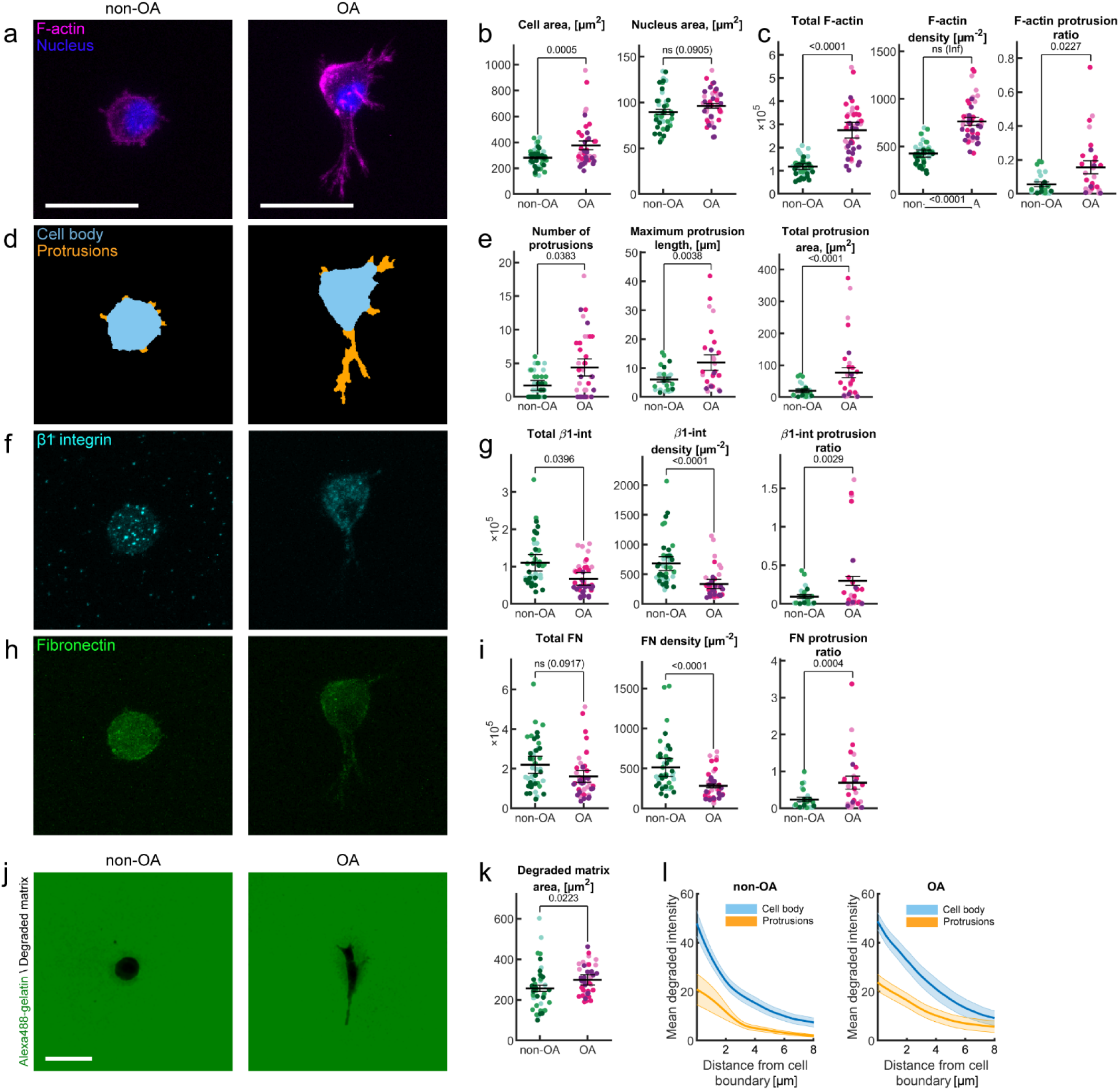
3D immunostaining analysis of cytoskeletal organization, cell-matrix adhesion and matrix degradation in non-OA and OA chondrocytes. **a** Maximum intensity projection of F-actin expression (magenta) and nucleus (blue) for a representative non-OA and OA chondrocyte. **b** Cell area and nucleus area. **c** Quantification of F-actin expression: Total F-actin intensity for a cell, F-actin density (computed as total F-actin intensity normalized by cell area), F-actin protrusion ratio (computed as F-actin intensity in the protrusions divided by F-actin intensity in the cell body). **d** Automatic segmentation of cell body (light blue) and protrusions (orange) for corresponding non-OA and OA chondrocyte. **e** Protrusion quantification (per cell): Number of protrusions, Maximum protrusion length, Total protrusion area. **f** Maximum intensity projection of β1 integrin expression (cyan) for a representative non-OA and OA chondrocyte. **g** Quantification of β1 integrin expression: Total β1 integrin intensity for a cell, β1 integrin density (computed as total β1 integrin intensity normalized by cell area) and β1 integrin protrusion ratio (computed as β1 integrin intensity in the protrusions divided by β1 integrin intensity in the cell body). **h** Maximum intensity projection of fibronectin (FN) expression (green) for a representative non-OA and OA chondrocyte. **i** Quantification of fibronectin expression: Total fibronectin intensity for a cell, Fibronectin density (computed as total fibronectin intensity normalized by cell area), Fibronectin protrusion ratio (computed as Fibronectin intensity in the protrusions divided by fibronectin intensity in the cell body). **j** Degradation assay: minimum intensity projections of fluorescently labeled gelatin (green) degraded by a representative non-OA and OA chondrocyte. **k** Quantification of total degraded matrix area using the degradation assay. **l** Mean degradation around cell body (light blue) and protrusions (orange) for increasing distance away from cell boundary (in µm). All statistical comparisons were performed using generalized linear mixed-effects models to account for the nested structure of the data. Error bars represent SEM, scalebars are 30 µm. Each green and magenta hue in the plots corresponds to a different patient (N=3 for each condition), each dot represents a single cell measurement (n≥15 for every patient).

Despite their larger size and more protrusive phenotype, OA chondrocytes exhibited significantly decreased total β1 integrin levels and, consistent with the trend observed in 2D, decreased β1 integrin density compared to non-OA cells (Fig. 3f, 3g). At the same time, OA chondrocytes displayed a higher β1 integrin localization in protrusions relative to the cell body (Fig. 3g, right). Total fibronectin deposition was significantly lower for OA chondrocytes, both in terms of total amount and density (Fig. 3h and Fig. 3i, left and middle), which was different from 2D findings, where OA led to higher fibronectin amount (but not density). Consistent with the increased β1 integrin localization in protrusions, we also observed a higher fibronectin localization in protrusions for OA chondrocytes (Fig. 3i, right). Apart from fibronectin deposition, we also assessed hydrogel degradation to have a more complete picture of ECM remodeling. Degradation was analyzed using a degradation assay where chondrocytes were embedded in a 3D PEG hydrogel mixed with an Alexa488 dye labeled gelatin to obtain a hydrogel with uniform fluorescence intensity in the absence of degradation (Fig. 3j). Quantification of the degraded area (including the area occupied by the cell) revealed that OA chondrocytes demonstrate a higher total hydrogel degradation (Fig. 3k) in line with their larger cell area. While total degraded area was found to be larger for OA chondrocytes, the width of the pericellular hydrogel region affected by degradation did not differ for OA versus non-OA chondrocytes (Fig. S8a). Within this degradation-affected region we also observed that degradation occurred primarily around the cell body rather than at protrusions for both OA and non-OA cells, as can be deduced from the stronger decrease in fluorescence intensity around the cell body compared to protrusions (Fig. 3l and Fig. S7). Degradation patterns around cell body and protrusions were not affected by OA, with OA and non-OA cells displaying comparable fluorescence intensity values as a function of distance from the cell border (Fig. 3l) and similar mean degradation curve profiles (Fig. S8b).

Collectively, these findings illustrate that OA chondrocytes cultured in a 3D degradable PEG-RGD hydrogel environment exhibit distinct morphological, cytoskeletal, adhesional and matrix alterations—namely, increased cell volume, enhanced actin polymerization, more pronounced protrusive phenotype, reduced expression of β1 integrin and fibronectin and relocalization of molecules of interest towards protrusions—compared to non-OA cells.

### 4. OA chondrocytes demonstrate enhanced force generation capacity in 3D degradable hydrogels

To test whether the observed protrusive phenotype and associated molecular changes (including relocalization) enhance force generation of OA chondrocytes, we performed 3D TFM experiments using the same PEG hydrogel system, in which 200 nm fluorescent beads were embedded (Fig. 4a). Cells were encapsulated in the gel and imaged 24 hours after seeding (Fig. 4b). OA chondrocytes induced significantly higher pulling (inward oriented) matrix displacements and exerted higher pulling tractions than non-OA chondrocytes (Fig. 4c). The more contractile phenotype of OA chondrocytes was also apparent from the larger fraction of pulling tractions per cell (Fig. 4d, left: 0.51±0.02 on average for OA, and 0.35±0.07 for non-OA) and a higher value of the global cell contractility metric (Fig. 4d, right: 54±48 Pa on average for OA, and −295±140 Pa for non-OA; positive and negative sign correspond to contractile and extensile activity respectively). Interestingly, both OA and non-OA chondrocytes also exhibited substantial pushing activity (i.e. outward oriented matrix displacements and tractions) and an individual chondrocyte (regardless of being OA or non-OA) typically displayed distinct areas of both pushing and pulling activity at the same time (Fig. S10). For non-OA chondrocytes pushing and not pulling was even governing the average force exerting behavior, as one can deduce from the average values of the fraction of pulling tractions (less than 0.5 for non-OA chondrocytes, while higher than 0.5 for OA chondrocytes) and the global contractility metric (negative for non-OA chondrocytes, while positive for OA chondrocytes). Further quantitative analysis of the pushing displacements and tractions demonstrated that they were of comparable order of magnitude to the pulling displacements and tractions for both OA and non-OA chondrocytes and that their magnitude was not affected by OA (Fig. 4e). While pulling tractions were located solely at the cell protrusions, pushing tractions were detected near the cell body and related to an increase in cell volume (or reversely, a decrease in cell volume upon addition of Cytochalasin D for obtaining a relaxed state) (Fig. S3).

**Figure 4.**
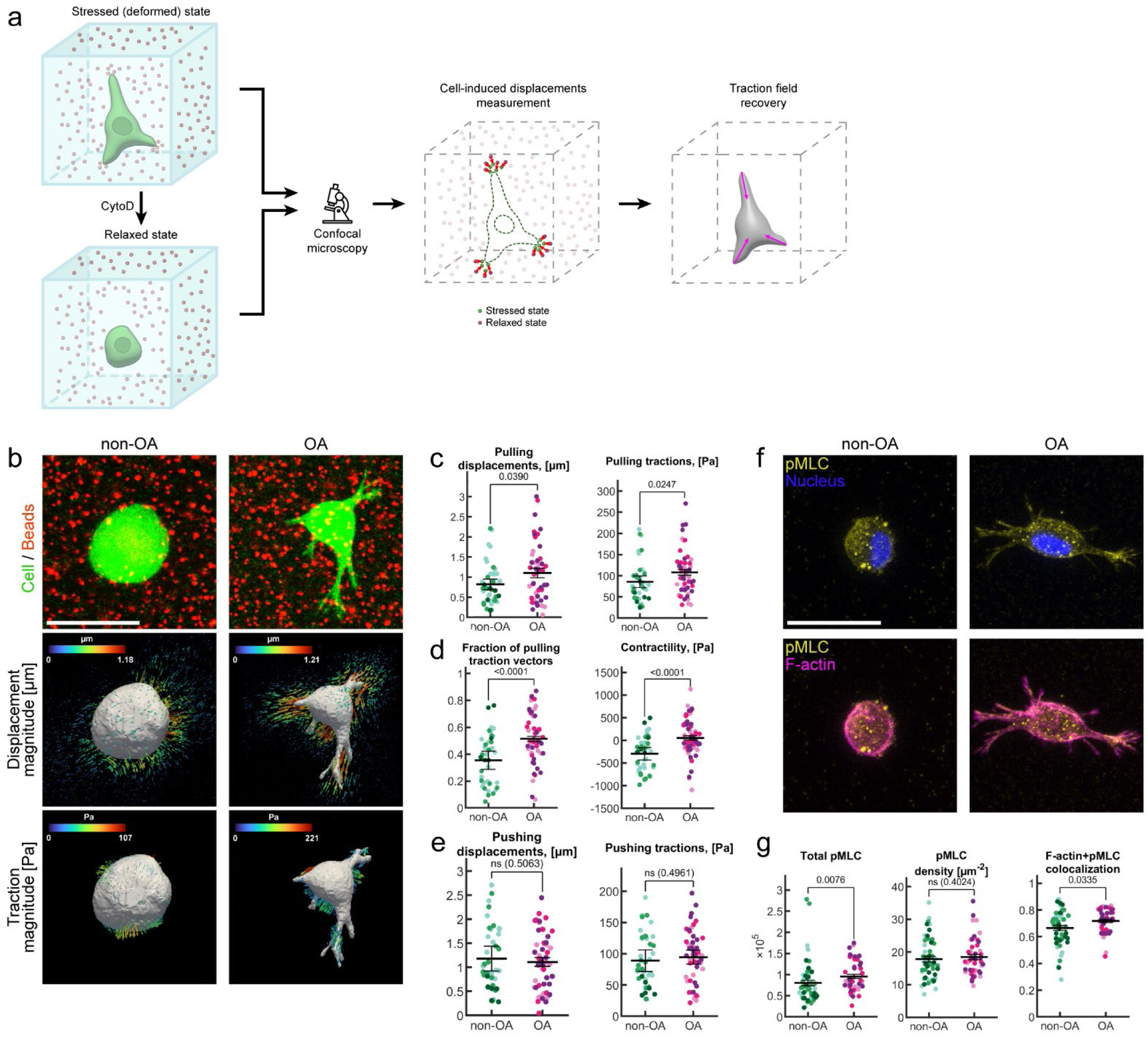
3D TFM analysis of cell tractions and contractility in OA and non-OA chondrocytes. **a** 3D TFM workflow. **b** Representative images illustrating the TFM workflow. (top) Representative maximum projected images of beads (red) and chondrocyte (green) for one non-OA and one OA chondrocyte. (middle) Corresponding 3D displacement magnitude renders. (bottom) Corresponding 3D traction magnitude renders. **c** Quantification of the 99^th^ percentile pulling (inward oriented) bead displacement magnitude [µm] and pulling (inward oriented) traction magnitude [Pa]. **d** Fraction of pulling traction vectors (with respect to total number of traction vectors) per cell. Contractility, a global metric of a cell’s ability to generate contractile forces, computed as the sum of all (signed) traction vector components (positive if tensile, negative if compressive) along the axis that connects the corresponding node to the cell centroid. **e** Quantification of 99^th^ percentile pushing displacements [µm] and 99^th^ percentile pushing tractions [Pa]. **f** Maximum intensity projection of pMLCexpression (yellow) and overlay with F-actin (magenta) for a representative non-OA and OA chondrocyte. **g** Quantification of pMLC expression: total pMLC intensity of a cell, pMLC density (computed as the ratio of total pMLC intensity normalized by the cell area). Colocalization of F-actin with pMLC. All statistical comparisons were performed using generalized linear mixed-effects models to account for the nested structure of the data. Error bars represent SEM, scalebars are 30 µm. Each green and magenta hue in the plots corresponds to a different patient (N=3 for each condition), each dot represents a single cell measurement (n≥15 for every patient).

Immunostaining showed increased pMLC content in OA chondrocytes (Fig. 4f, Fig. 4g, left). This increase in pMLC was proportional to the observed increase in cell area for OA chondrocytes, resulting in similar pMLC densities for OA and non-OA cells (Fig. 4g, middle), which is in agreement with our previous 2D observations. However, opposite to 2D, we also observed increased colocalization of F-actin and pMLC in OA chondrocytes (Fig. 4g, right), and together with the elevated F-actin levels mentioned earlier (Fig. 3c) it suggests stronger actomyosin coupling in OA, consistent with the higher pulling tractions and enhanced contractility of OA chondrocytes (Fig. 4c, right, and Fig. 4d, right).

## Discussion

Recent advances in OA research have emphasized the importance of mechanical loading in disease progression, primarily by studying chondrocyte response to external forces [28], [29]. However, intrinsic chondrocyte-generated forces and the molecular machinery underlying force exertion remain poorly understood, particularly in physiologically relevant 3D microenvironments. Our study addressed these gaps by investigating how OA affects chondrocyte force exertion and how culture dimensionality influences these mechanical phenotypes.

A key finding of our study is that OA-affected chondrocytes display a distinct contractile and protrusive phenotype—but only when embedded in a 3D degradable matrix, revealing a disease-specific mechanical behavior that cannot be captured by conventional 2D cultures. This protrusive phenotype aligns with previous *in vivo* observations of enhanced protrusive morphologies in chondrocytes in human OA cartilage, particularly in regions of cartilage degeneration [30], [31], [32], [33], [34]. The ability of our 3D model to recapitulate this protrusive behavior supports its physiological relevance. Importantly, we extend these findings by showing that OA chondrocytes also exert higher contractile forces in 3D, which colocalize at protrusion tips. This suggests a direct link between protrusive activity and contractility.

In contrast, 2D non-degradable hydrogel cultures failed to reveal any OA-associated differences in force exertion or protrusion formation. To investigate whether OA nonetheless altered the molecular force machinery in 2D, we assessed the expression and localization of cytoskeletal (F-actin, pMLC) and proteins involved in cell-ECM adhesion (β1 integrin and fibronectin). OA chondrocytes showed decreased cortical localization of all cytoskeletal and adhesion proteins and increased expression only for pMLC. Although pMLC is commonly regarded as a driver of actomyosin contractility, increased levels alone did not lead to higher force output in 2D.

In 3D, however, the OA-induced changes in cytoskeletal organization were more pronounced and led to elevated force generation. While total pMLC levels were similarly increased in both 2D and 3D, OA chondrocytes in 3D also exhibited increased levels of F-actin and enhanced colocalization of F-actin and pMLC. In addition, OA was also associated with a shift of F-actin, β1 integrin and fibronectin towards protrusion areas (relative to the cell body), which could all have contributed to more effective force transmission to the ECM. Apart from the shift in β1 integrin localization, we observed an overall decrease in β1 integrin in OA chondrocytes in our 3D system (unlike in 2D), contrary to a previous study that observed no difference in β1 integrin or α5β1 for OA cartilage explants [13], [14]. However, another study has linked reduced β1 integrin levels to more severe cartilage damage [15] in line with the results observed in our degradable hydrogel system.

One limitation of our study is the extent to which our 3D hydrogel model replicates the native cartilage ECM, as well as our inability to decouple the individual effects of matrix degradability and culture dimensionality. While our degradable hydrogels incorporate OA-relevant features— such as MMP-cleavable crosslinks (responsive to MMP1 and MMP3, which are upregulated in OA) and RGD peptides supporting fibronectin-mediated β1 integrin binding—their stiffness (Young’s modulus ∼520 Pa) is substantially lower than that of native cartilage (e.g., PCM: ∼137 kPa; OA: ∼96 kPa [35]). However, this reduced stiffness is necessary to enable sufficiently large cell-generated hydrogel deformations that can be measured using TFM. The softer hydrogel may also accelerate degradation and protrusive behavior relative to native tissue, allowing us to observe OA-specific phenotypes over a short (24-hour) culture period. Despite these limitations, our 3D degradable hydrogel serves as a physiologically relevant and reductionist model that recapitulates hallmark features of OA pathology—such as increased protrusive activity and degradation—while enabling the quantification of chondrocyte-generated forces through TFM.

An important consideration when combining TFM with degradable hydrogels is the potential impact of matrix degradation on traction calculations, as degradation can lead to local hydrogel softening that is not accounted for in the calculations. To ensure valid comparisons of force generation between OA and non-OA chondrocytes, we demonstrated that patterns of degradation intensity were the same for OA and non-OA chondrocytes, both around protrusions and around the cell body. This supports the reliability and interpretability of our traction calculations. In addition, the degradation assay also enabled to assess local differences in degradation intensity: degradation was more prominent around the cell body than at the protrusions, which was not affected by OA, and which is in line with previous observations in cancer cells and endothelial cells [36]. The cell body was typically associated with regions of pushing forces, which in turn could be linked to cell volumetric growth. Similar phenomena have been reported in confined environments, where pushing forces arise as cells expand within spatial constraints [37]. Such pushing forces may cooperate with MMP-mediated ECM degradation to create space for volumetric growth, following similar cooperation proposed for cancer cell invadopodia [38]. The magnitude of chondrocyte pushing forces was similar across OA and non-OA conditions. While pushing forces are typically lower in magnitude than pulling forces as previously reported for mammary epithelial cells [39], they appeared comparable for chondrocytes—likely due to the overall low traction levels exerted by chondrocytes (pulling tractions of 80 Pa in chondrocytes versus ∼150 Pa in HUVECs)[40].

Together, our findings reveal a distinctive contractile and protrusive phenotype in OA chondrocytes within 3D degradable matrices. This combination of enhanced contractility, protrusive activity, and matrix degradation may contribute to several hallmark features of OA pathology. These phenotypes are reminiscent of migratory behavior seen in other disease contexts, such as cancer and vascular malformations, where protrusive forces and matrix degradation facilitate invasion [40], [41]. In OA cartilage, such features may underlie chondrocyte clustering and lacunae formation—particularly in degenerated cartilage regions—where migration is thought to drive aggregation due to the increased presence of protrusions [42].

Mechanistically, our data could support the idea that elevated intrinsic contractility may also be a driver of matrix degradation. Prior studies have shown that inhibition of contractility-related pathways (e.g., ROCK, myosin II, or actin polymerization) reduce MMP expression in chondrocytes and cartilage breakdown [16], [43]. In contrast, the enhanced contractility observed in our study may inversely actively sustain matrix degradation. Moreover, protrusive cells in OA cartilage show increased levels of cytokine IL-1β, which are known to initiate MMP signaling and cartilage degradation [44]. Taken together, this suggests a mechanochemical feedback loop in which force-generating, protrusive cells promote local cytokine signaling and MMP-mediated ECM degradation. Finally, OA-associated alterations in chondrocyte force generation and force generating machinery (as observed here for F-actin, pMLC, β1 integrins, fibronectin) may impact how chondrocytes sense, transmit and respond to external mechanical loading. In conclusion, it makes disrupted chondrocyte force generation a promising new target in the study of OA pathogenesis.

## Methods

### 1. Cell culture

Non-OA and OA primary cells were isolated from cartilage of patients that underwent hip replacement surgery in the university hospital Gasthuisberg, KU Leuven. Non-OA primary cells were collected from three patients that had surgery for osteoporotic or malignancy associated fractures with no history of OA (two women, one man, age 44, 77 and 90). OA primary cells were obtained from three OA patients undergoing hip surgery (one woman, two men, age 64, 71 and 82). Primary human articular chondrocytes (hACs) were isolated as previously described [28]. Briefly, cartilage was first dissected from the joint surface, rinsed with phosphate buffered saline (PBS), and cut into small pieces. The cartilage pieces were incubated with 2 mg/ml pronase solution (Roche) for 90 minutes at 37°C and digested overnight at 37°C in 1.5 mg/ml collagenase B solution (Roche). Then, the preparation was filtered through a 70 µM strainer and cells were plated in culture flasks (T75, Avantor) and cultured in a humidified atmosphere at 37°C and 5% CO_2_. Culture medium consisted of Dulbecco’s Modified Eagle Medium F12 (DMEM/F12, Gibco), 10% fetal bovine serum (FBS, Gibco), 1% (vol/vol antibiotic/antimycotic (Gibco), and 1% L-glutamine (Gibco). All cells were used between P0 and P2.PAA hydrogel preparation Polyacrylamide (PAA) hydrogels with a Young’s modulus of 4.5 kPa were prepared according to previously reported protocol [45]. Fluorescent beads (Invitrogen, FluoSpheres Carboxylate-Modified Microspheres, 200 nm) were added to the hydrogel mixture (1:60 concentration) for TFM samples. Briefly, MilliQ, sonicated fluorescent beads, 40% Acrylamide (Sigma-Aldrich, A4058) and 2% bisacrylamide (Sigma-Aldrich, M1533) were mixed in that order. Hydrogel polymerization was initialized by ammonium persulfate (4.4 µM) and catalyzed by N,N,N’,N’-tetramethylethylenediamine (Sigma-Aldrich, T7024). A 15 µl hydrogel droplet was placed on the bottom of a 6 well plate with glass bottom (Cellvis, P06-14-0-N), pre-treated with bind-silane (Serva, 28739.01). The hydrogel was flattened with a coverslip glass, pre-treated with Sigmacote and left to polymerize at room temperature for 30 minutes. The gel surface is functionalized by two rounds of 0.67 µM Sulfo-SANPAH (Sigma-Aldrich, 803332) treatment under UV-irradiation with 365 nm wavelength emission for 30 minutes, followed by a 2 hour incubation at 37°C in 10 µg/ml Fibronectin (human plasma, Sigma-Aldrich) diluted in 0.1% v/v acetic acid at pH 3.4 for homogeneous coating [46]. 3000 cells were seeded per hydrogel and allowed to spread for 24 hours. Afterwards, samples were either live imaged for TFM experiments or fixed for half an hour in 4% paraformaldehyde for immunostaining. PAA hydrogels were mechanically characterized as previously described in [47] and assuming a Poisson’s ratio of 0.49.

### 2. PEG hydrogel preparation

Degradable polyethylene glycol (PEG) precursors and buffers were prepared as previously described [40]. Fluorescent beads (Invitrogen, FluoSpheres Carboxylate-Modified Microspheres, 200 nm) were added to the hydrogel mixture (1:60 v/v concentration) for TFM samples. Briefly, milliQ water, sonicated fluorescent beads, 10X Buffer, lysine-RGD (50 µM, Pepmic), PEG and cell solution were mixed in that order. The cell concentration was calculated to have 3000 chondrocytes per hydrogel droplet. Enzymatic crosslinking was initialized by addition of activated FXIII. We used a w/v hydrogel composition of 1.2% that corresponded to a Young’s modulus of 520 Pa, as mechanically characterized previously [40] and assuming a Poisson’s ratio of 0.3. An 18 µl hydrogel droplet was placed on the bottom of an Imaging chamber (µ-Slide 18 Well, Ibidi) and allowed to crosslink at 37°C and 5% CO_2_ for 30 minutes. Afterwards culture medium was added, and cells were incubated for 24 hours. Afterwards, samples were either live imaged for TFM experiments or fixed for half an hour in 4% paraformaldehyde for immunostaining and matrix degradation quantification.

### 3. 2D Immunostaining

Hydrogels for immunostaining were fixed for half an hour in 4% paraformaldehyde followed by washing with PBS.

Immunostaining of pMLC requires permeabilization of the cell membrane with Permeabilization solution constituting 0.3% Triton X (10%) in PBS 1X at room temperature for 1 hour. Subsequently, samples were treated with permeabilizing blocking solution constituting 1% BSA, 10% goat serum and 1% Triton X (10%) at room temperature for four and half hours. Primary antibody Phospho-Myosin Light Chain 2 (Ser19) (Cell Signaling Technology #3671 1:100) was suspended in permeabilizing blocking solution and applied to the sample and incubated at 37°C and 5% CO_2_ for 1 hour and half. The sample was washed with rinsing solution constituting 0.05% TWEEN 20 in PBS 1X and incubated at 4°C on a shaker overnight. The following day, three washes with rinsing solution were carried out every two hours. Goat anti-Rabbit IgG secondary antibody conjugated with Alexa Fluor 488 (Invitrogen, 1:1000) was diluted in permeabilizing blocking solution and incubated at 37°C and 5% CO_2_ for 1 hour and half. Afterwards, the sample was washed with rinsing solution and put overnight on a shaker at 4°C. The following day, three washes with rinsing solution were carried out every two hours.

For immunostaining of β1 integrin and fibronectin, the following protocol was followed. First, samples were treated with non-permeabilizing blocking solution constituting 1% BSA, 10% goat serum and 0.05% Tween 20 for four hours and half. Primary antibodies were suspended in non-permeabilizing blocking solution and applied to the sample and incubated at 37°C and 5% CO_2_ for 1 hour and half. Anti-fibronectin antibody (Sigma-Aldrich, F3648 1:1000) and anti-β1 Integrin antibody (BD Biosciences #553715, 1:200) were used as primary antibodies. The following day, three washes with rinsing solution were carried out every two hours and the sample was kept at 4°C on a shaker in between washes. Goat anti-Rabbit IgG secondary antibody conjugated with Alexa Fluor 488 (Invitrogen, 1:1000) and Goat anti-Rat IgG secondary antibody conjugated with Alexa Fluor 546 (Invitrogen, 1:1000) were diluted in non-permeabilizing blocking solution and incubated at 37°C and 5% CO_2_ for 1 hour and half. Afterwards, the sample was washed with rinsing solution and incubated at 4°C on a shaker overnight. The following day, three washes with rinsing solution were carried out every two hours and the sample was kept at 4°C on a shaker in between washes.

Then, for all 2D samples, phalloidin conjugates with Atto 647 (Sigma-Aldrich, 1:2000) were diluted in PBS and applied to the sample, incubating on a shaker at 4°C for 45 minutes. Finally, DAPI (1:5000) diluted in PBS was applied to the sample and incubated for ten minutes at room temperature. Samples were extensively washed with rinsing solution before imaging.

### 4. 3D Immunostaining

Hydrogels for immunostaining were fixed for half an hour in 4% paraformaldehyde followed by washing with PBS.

Immunostaining of pMLC requires permeabilization of the cell membrane with Permeabilization solution constituting 0.3% Triton X (10%) in PBS 1X at 37°C and 5% CO_2_ for half an hour. Subsequently, samples were treated with permeabilizing blocking solution constituting 1% BSA, 10% goat serum and 1% Triton X (10%) at room temperature for four hours and half. Primary antibody Phospho-Myosin Light Chain 2 (Ser19) (Cell Signaling Technology #3671 1:50) was suspended in permeabilizing blocking solution and applied to the sample and incubated at 4°C on a shaker overnight. The following day, five washes with rinsing solution were carried out every two hours and the sample was kept at 4°C on a shaker in between washes. Then, Goat anti-Rabbit IgG secondary antibody conjugated with Alexa Fluor 488 (Invitrogen, 1:500) was diluted in permeabilizing blocking solution and incubated at 4°C on a shaker overnight. The following day, five washes with rinsing solution were carried out every two hours and the sample was kept at 4°C on a shaker in between washes.

For immunostaining of β1 integrin and fibronectin, the following protocol was followed. First, samples were treated with non-permeabilizing blocking solution constituting 1% BSA, 10% goat serum and 0.05% Tween 20 for four hours and half. Primary antibodies were suspended in non-permeabilizing blocking solution and applied to the sample and incubated at 4°C on a shaker overnight. Anti-fibronectin antibody (Sigma-Aldrich, F3648 1:500) and anti-β1 Integrin antibody (BD Biosciences #553715, 1:100) were used as primary antibodies. The following day, five washes with rinsing solution were carried out every two hours and the sample was kept at 4°C on a shaker in between washes. Goat anti-Rabbit IgG secondary antibody conjugated with Alexa Fluor 488 (Invitrogen, 1:500) and Goat anti-Rat IgG secondary antibody conjugated with Alexa Fluor 546 (Invitrogen, 1:500) were diluted in non-permeabilizing blocking solution and incubated at 4°C on a shaker overnight. The following day, five washes with rinsing solution were carried out every two hours and the sample was kept at 4°C on a shaker in between washes.

Then, for all 3D samples, phalloidin conjugates with Atto 647 (Sigma-Aldrich, 1:1000) were diluted in PBS and applied to the sample, incubating on a shaker at 4°C for four hours. Finally, DAPI (1:2000) diluted in PBS was applied to the sample and incubated for ten minutes at room temperature. Samples were extensively washed with rinsing solution before imaging.

### 5. Traction force microscopy image acquisition

All live cell imaging for TFM was done using a Leica TCS SP8 confocal fluorescence microscope. For TFM, confocal images were acquired using a 40X water objective using 1.5 zoom at a 512×512 px resolution at 1AU and bidirectional imaging with phase correction. Excitation wavelengths were set to 488 nm (green cell channel) and 563 nm (red beads channel). The detection wavelengths of the hybrid detectors were set to 495-555 nm and 570-630 nm respectively to minimize crosstalk between the green and red channel and allow for simultaneous imaging. Stressed state images were collected as z-stacks with 0.5 µm step size (final voxel size of 0.38×0.38×0.5 µm).

For 2D samples, relaxed state was induced by removing the cells by addition of a droplet of 5% Sodium dodecyl sulphate (SDS). After 15 minutes, the hydrogel returned to a stress-free state and relaxed state bead images were acquired using the same imaging settings.

For 3D samples, relaxed state was induced by depolymerizing actin by addition of Cytochalasin D (dissolved in DMSO, Sigma Aldric) at a final concentration of 4 µM. After 1 hour, the hydrogel returned to a stress-free state and relaxed state bead images were acquired using the same imaging settings.

### 6. 2D traction force microscopy analysis

2D TFM analysis was performed using a previously described computational framework implemented in MATLAB [47]. Briefly, beads and cell images were filtered using a Gaussian kernel to suppress noise. Cell masks, used to calculate the metrics in Fig. 2, were obtained by thresholding the cell channel. Unwanted spatial shifts between the stressed and relaxed state image stacks were corrected for by rigid image registration using the *findshift* function from the DipImage library [48]. Cell-induced displacements in the bead images were measured using a non-rigid, free-form deformation image registration method [49] using the Elastix toolbox [50]. Tractions were recovered by considering PAA gels as homogeneous, isotropic, linear elastic half spaces and applying a modified Tikhonov regularized Fourier transform traction cytometry algorithm as previously described [45], [51]. Since noise levels in the measured displacement fields were similar throughout the dataset, a constant value of 10 was set for the Tikhonov regularization parameter lambda. 99^th^ percentile displacements, 99^th^ percentile tractions (both referred to as maximum displacement and tractions in the text), contractility and cortical traction ratio were restricted to the cell mask. The boundary of the cell mask was obtained and then dilated inwards with a disk-shaped structuring element of an equivalent physical size of around 5 µm (see Fig. S5). The resultant region was considered as the cell cortex. The cortical traction ratio is computed as the ratio between the sum of traction magnitudes inside the cell cortex over the sum of all traction magnitudes within the cell mask. The contractility of each cell is defined by the following formula:

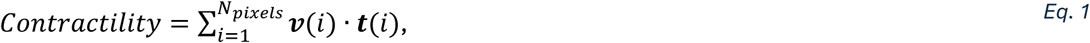

where *N_pixels_* is the number of pixels within the cell mask, *v*(*i*) is the unit vector that points from pixel *i* to the centroid of the mask, *t*(*i*) is the traction vector at pixel *i*, and · is the dot product (see also Fig. S9).

### 7. 3D traction force microscopy analysis

3D TFM images were analyzed using the TFMLAB software toolbox (available on: https://gitlab.kuleuven.be/MAtrix/Jorge/tfmlab_public) [52] and as described in [40]. The hydrogel was modeled as a linear elastic material with Young’s Modulus E=520 Pa and Poisson ratio ν = 0.3. TFMLAB computes tractions using a physics-based inverse method that is implemented in a finite element framework and that looks for a displacement field similar to the measured one while fulfilling equilibrium of forces in the hydrogel domain [53], [54]. To analyze the TFM data, different metrics were defined. Cells were considered ‘active’ if more than 1% of the voxels in their vicinity had a displacement magnitude larger than the pixel size (0.38 µm). A cell is deemed passive otherwise and discarded since displacements are noise-level. To determine whether a traction is pulling or pushing, the traction vector at each finite element node was projected onto the vector that connects the node with the centroid of the cell mesh (*v*(*i*) · *t*(*i*) in). Then, based on the sign of the projected traction, it can be categorized as pulling (positive sign) or pushing (negative sign). To avoid outliers, the ‘maximum’ pulling and pushing traction are computed using this categorization as the 99^th^ percentile traction (Fig. 4c). The fraction of pulling cells displayed in Fig. 4d is the fraction of positive traction vectors per cell. The contractility of each cell (Fig. 4d) is defined by taking the sum of the tractions projected on the connecting vectors using Eq. 1, where, instead of summing over the pixels, we sum over the finite element nodes of the cell surface mesh (see also Fig. S9).

### 8. Immunostaining image acquisition

All Immunostaining imaging was done using a Leica TCS SP8 confocal fluorescence microscope. Confocal images were acquired using a 25X water objective using 3.4 zoom at a 512×512 resolution at 1AU. Excitation wavelengths were set to 405 nm (Blue nucleus channel), 488 (Green pMLC or Fibronectin channel), 563 nm (Red β1 integrin channel) and 647 (Far-red F-actin channel). The emission wavelengths of the Hybrid detectors were 410-460 nm, 495-555 nm, 570-630 nm and 666-753 nm respectively. To avoid crosstalk between the channels, they were imaged sequentially: sequence 1 - Blue and Far-red channel, sequence 2 - Green channel and sequence 3 - Red channel. 3D image z-stacks were acquired with a 0.5 µm step size.

For imaging of the degradation assay, a Leica TCS SP8 confocal fluorescence microscope was used. Confocal images were acquired using a 25X water objective using 3.4 zoom at a 512×512 resolution at 1AU and bidirectional imaging with phase correction. Excitation wavelengths were set to 405 nm (Blue nucleus channel), 488 nm (Green gelatin channel) and 647 nm (Far-red F-actin channel). The emission wavelengths of the Hybrid detectors were set to were 410-460 nm, 495-555 nm and 670-730 nm respectively. To avoid crosstalk between the channels, they were imaged sequentially: sequence 1—Blue and Far-red channels and sequence 2—Green channel. 3D image z-stacks were acquired with a 0.5 µm step size (final voxel size of 0.27×0.27×0.5 µm).

### 9. Quantification of cell and nuclear morphology

3D fluorescence F-actin image stacks of 2D and 3D samples are maximally projected and filtered with a median filter. Cell segmentation of the cells was done using a manually adjusted intensity threshold to obtain a cell mask. Area, Solidity, Eccentricity, and Circularity data as shown in Fig. S1a and Fig. S2a were computed using the *regionprops* function in MATLAB (see Fig. S4 for the definition of metrics). Protrusions were segmented as described in [40]. A schematic overview is depicted in Fig. S6. First, an opening operation on the binary mask using a disk-shaped structuring element was performed. The resulting binary image was subtracted from the cell mask, and small objects were removed.

For 2D samples, protrusions were segmented using a structure element with a size ∼8 μm and a minimum area of ∼140 μm^2^. For 3D samples, protrusions were segmented in two steps, using a structure element with a size ∼2 μm and a minimum area of ∼5 μm^2^ in a first step, and a structure element with a size ∼5 μm and a minimum area of ∼25 μm^2^ in a second step. The number of protrusions were counted, and the area and maximum length was calculated using *regionprops*.

3D fluorescence DAPI image stacks of 2D and 3D samples are maximally projected and filtered with a median filter. Segmentation of the nucleus was done using automated Otsu thresholding. Nuclear area was computed using the *regionprops* function in MATLAB.

### 10. Quantification of actin

3D fluorescence F-actin image stacks of 2D and 3D samples are maximally projected. Total actin intensity metric was obtained by summing the actin intensity values within the cell mask. The actin density was obtained by normalizing the total actin intensity by the area of the cell mask. The cortical actin shown in Fig. 1d is calculated by taking the ratio of the sum of actin intensities within the cortex mask (obtained as described in Methods section 7) over the total actin intensity within the cell mask. F-actin protrusion ratio was computed as the ratio of total actin in the protrusions with respect to total actin in the cell body.

### 11. Quantification of pMLC and Fibronectin

3D fluorescence image stacks of 2D and 3D samples are maximally projected. Cell segmentation of the cells was done using a manually adjusted intensity threshold to obtain a cell mask. Total pMLC intensity was computed by summing the pMLC intensity within the cell mask. pMLC density was computed by normalizing the total pMLC intensity by the cell area. Total fibronectin was computed by summing the fibronectin intensity within the vicinity of the cell (computed with a dilation of the cell mask with a disk-shaped structure element of ∼2 μm), normalized to background signal. The fibronectin density is computed by normalizing the total fibronectin intensity by the cell area. For 2D samples, the cortical fibronectin and pMLC metric is computed as the ratio of intensity within the cortex over the total intensity. For 3D samples, fibronectin protrusion ratio shown in Fig. 3c is computed as the ratio of total fibronectin in the protrusion area with respect to total fibronectin in the cell body area (as defined in Fig. S6).

Colocalization of actin and pMLC (shown in Fig. 2e,right and Fig. 4g, right) within a cell is reported as the correlation coefficient and is computed using the projected fluorescent z-stacks and the *corr2* function in MATLAB. Prior to the computation, the respective fluorescent images are blurred using a Gaussian smoothing kernel with standard deviation of 0.5 (function *imgaussfilt* in MATLAB).

### 12. Quantification of β1 integrin

For the β1 integrin immunostaining, we were only interested in the β1 integrin located at the cell membrane in contact with the hydrogel, since this is where forces are transduced to the ECM. This required a different approach in 2D and in 3D samples.

In 2D samples we removed the top half part of the cell by segmenting the F-actin image with Otsu thresholding, locating the centroid of the resultant mask and discarding the image slices above this centroid. Then a maximum projection of the remaining 3D fluorescence β1 integrin image stack was done and used for the quantifications. Then, a maximum projection of the remaining 3D fluorescence β1 integrin image stack was done and used for the quantifications. The total β1 integrin intensity is computed by summing the β1 integrin intensity within the cell mask. The β1 integrin density is computed by normalizing the total β1 integrin intensity within the cell mask by the cell mask area. Cortical β1 integrin is computed as the ratio of total β1 integrin intensity within the cell cortex over the total β1 integrin intensity in the cell mask.

3D fluorescence β1 integrin image stacks of 3D samples are maximally projected. Cell segmentation of the cells was done using a manually adjusted intensity threshold to obtain a cell mask. A membrane mask was computed by combining the cell cortex and protrusions. The total β1 integrin intensity is computed by summing β1 integrin intensity within the membrane mask. The β1 integrin density is computed by normalizing the total β1 integrin intensity within the membrane mask by the membrane mask area. β1 integrin protrusion ratio was computed as the ratio of total β1 integrin intensity within protrusions over total β1 integrin intensity in the cell body (as defined in Fig. S6).

The colocalization of actin-β1 integrin within a cell is reported as the correlation coefficient and is computed as explained above using the projected fluorescent z-stacks. For 2D samples, correlation is computed within the cell mask. For 3D samples, correlation is computed within the membrane mask.

### 13. Quantification of matrix degradation

Gelatin labeled with Alexa488 dye (1:20 concentration) was added to the PEG hydrogel solution to give a uniformly fluorescent hydrogel. The hydrogel degraded area was quantified using a degradation assay as described in [40]. Briefly, the gelatin intensity image stack is minimum projected, inverted and binarized. Segmentation of the degraded hydrogel is used to compute the degraded matrix area (Fig. 3k). To distinguish the degradation around the cell body and the protrusions, the cell body mask and protrusion mask are dilated by one pixel in thirty steps and a mean degradation intensity within the dilated masks is computed for each step (Fig. S7). The mean degradation intensity curves for the cell body and protrusions displayed in Fig. 3l are computed using this workflow. Overall mean degradation around the protrusions is compared between OA and non-OA cells by averaging the stepwise mean degradation intensities for a perimeter of 5 μm (Fig. S8a). To evaluate the decay profile of the degradation curves, an exponential function is fitted on the cell body and protrusion degradation curve of each cell using the MATLAB function *fit* with the fittype ‘exp1’ (Fig. S8b).

### 14. Statistics

Statistical analysis was performed using generalized linear mixed-effects models (GLMMs) to account for the nested structure of the data (cells nested within patients, with patients grouped by two experimental conditions, namely OA and non-OA) and to test for differences between experimental conditions [55]. The choice of distribution and link function was tailored to the properties of each metric. For data that passed the Shapiro-Wilk normality test, a normal distribution was used. Non-normal, positive, unbounded data were modeled using a gamma distribution with a log link, while count data were analyzed using a Poisson distribution with a log link. For metrics bounded between 0 and 1, a logit transformation was applied, and the transformed data were modeled using a normal distribution. Model diagnostics were performed to ensure numerical stability and appropriate fit. Statistical significance was assessed using the Wald test. All analyses were conducted using MATLAB (version R2024b, MathWorks). This approach allows for robust statistical inference while accounting for the complex structure of the experimental data, as recommended by [56], [57].

## Acknowledgements

The authors acknowledge financial support from KU Leuven grant IDN/20/019 and iBOF/21/083 C. JBF is supported by an FWO postdoctoral fellowship 1259223N.

