## Supplementary material for "Osteoarthritic chondrocytes exert higher contractile forces and exhibit enhanced protrusive activity when cultured in 3D degradable hydrogels"

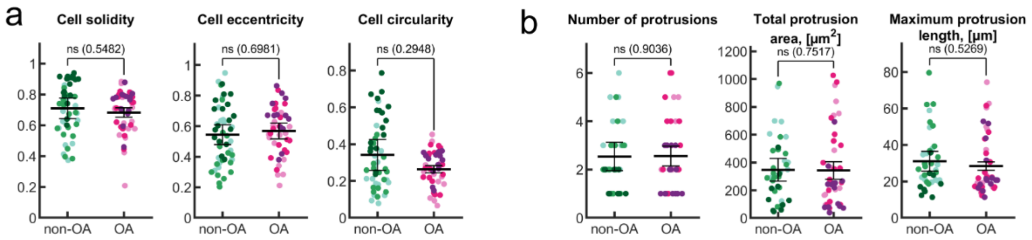


*Figure S1. **a** Quantification of cell morphology metrics for non-OA and OA chondrocytes cultured on 2D PAA substrates. **b** Quantification of protrusion metrics for non-OA and OA chondrocytes cultured on 2D PAA substrates.*

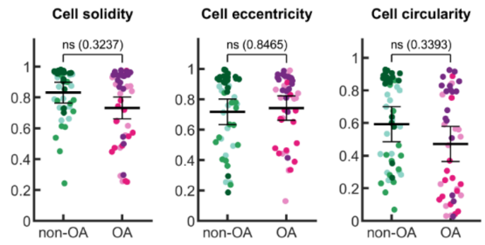


*Figure S2. Quantification of cell morphology metrics for non-OA and OA chondrocytes embedded in 3D degradable PEG hydrogels.*

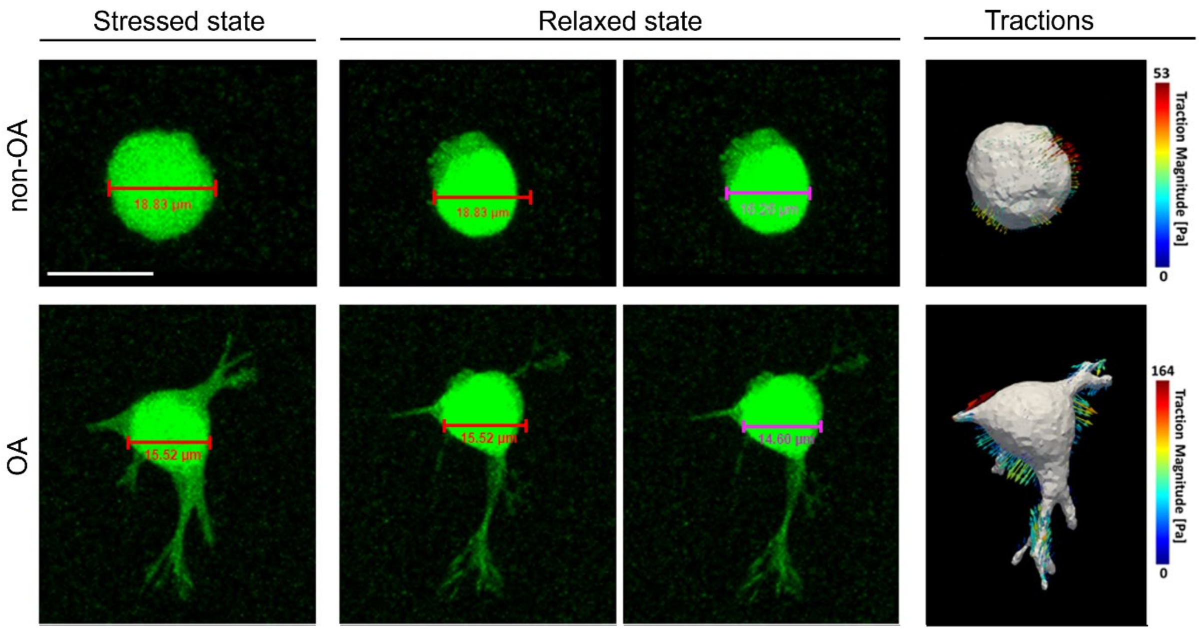


*Figure S3. Cell volume analysis before (stressed state) and after addition of Cyto D (relaxed state). Red scalebar indicates cell width in stressed state and purple scalebar indicates cell width in relaxed state. Cell width decrease after Cyto D application spatially correlated with location of pushing tractions. White scalebar is 20 μm.*

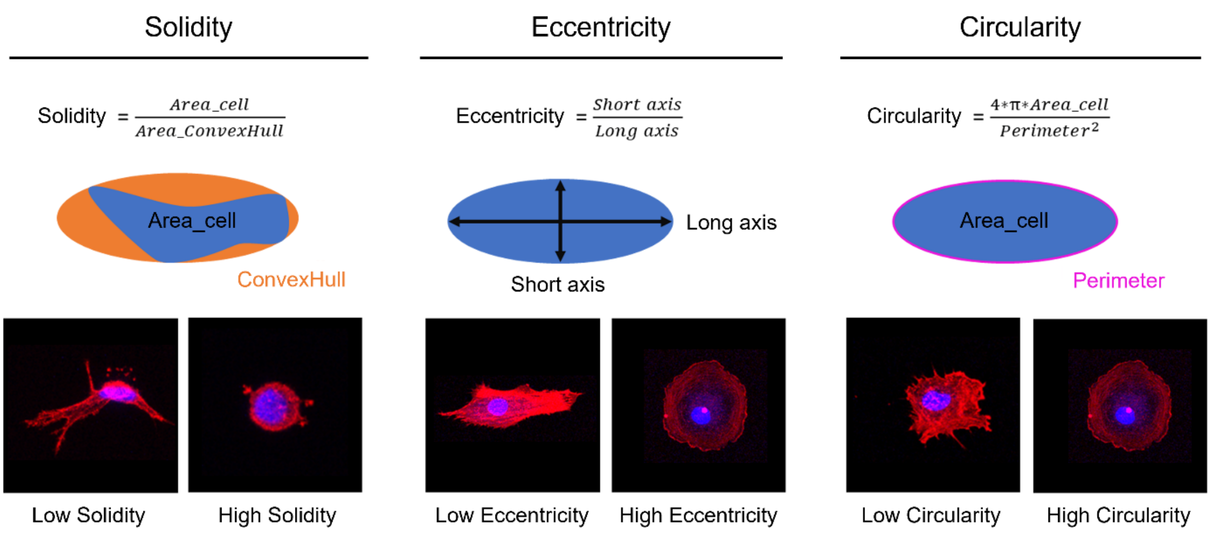


*Figure S4. Cell morphology metrics with cell indicative of a low or high value. Cell Solidity is computed as the ratio of the cell area and the area of the convex hull around the cell. Cell Eccentricity is computed as the ratio of the short axis and the long axis of the cell mask. Cell Circularity is computed as the cell area over the cell perimeter squared, multiplied by a factor of 4 times π.*

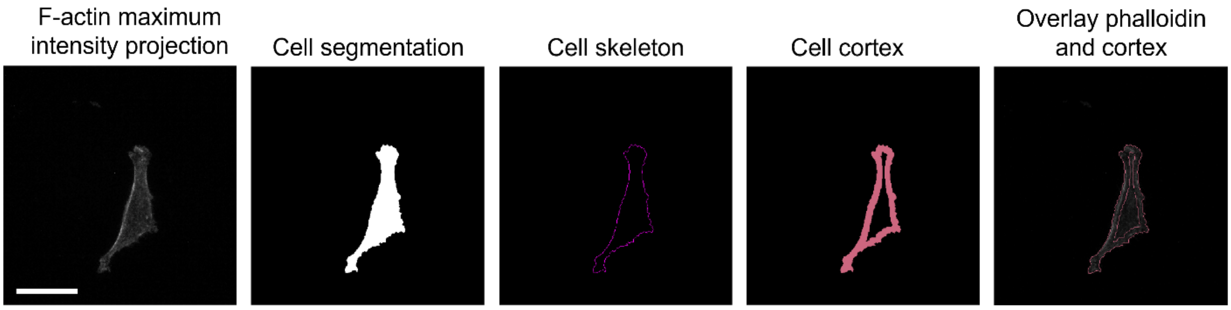


*Figure S5. Representative steps for cell cortex segmentation method. First, the F-actin intensity z-stack is maximum projected, and a cell mask is created using manual thresholding. From the cell mask, the outer cell boundary is selected as cell skeleton. Subsequently, the cell cortex is segmented by dilating the cell skeleton inwards with a disk-shaped structure element of size ~5 µm. The final step shows the overlay of the F-actin image and cell cortex. Scalebar is 30 µm.*

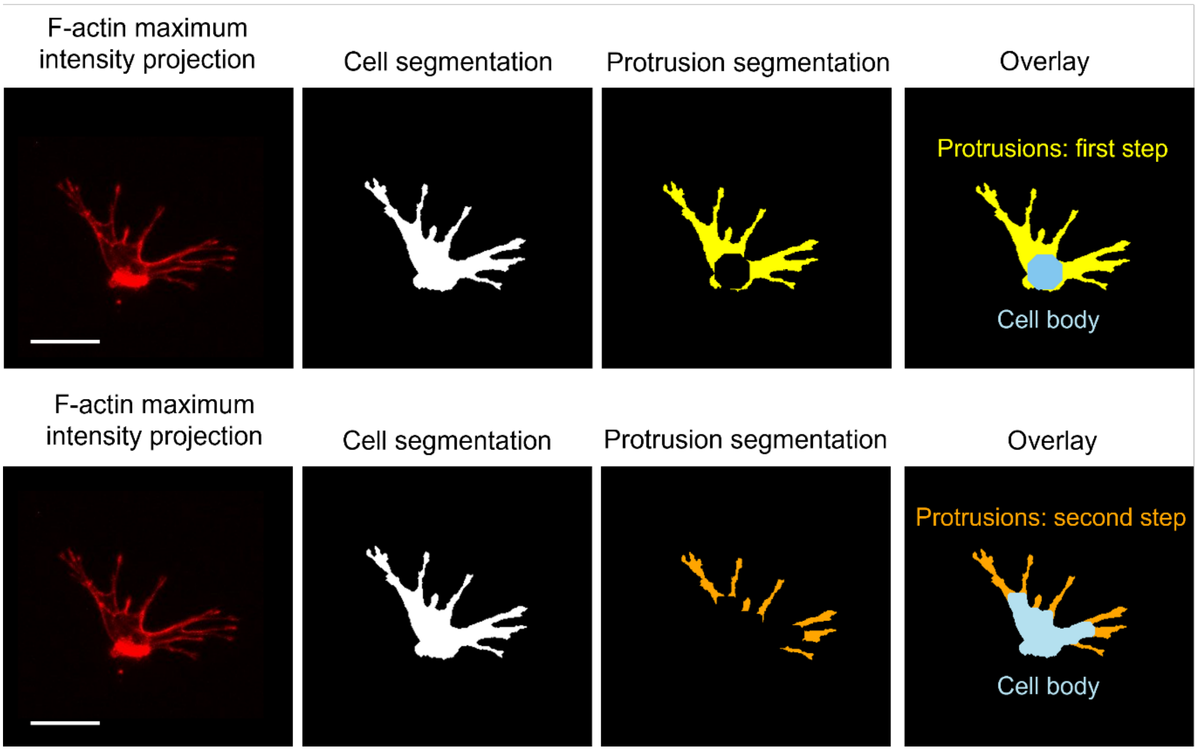


*Figure S6. Representative steps of protrusion segmentation method for cells in 3D. First, the F-actin intensity z-stack is maximum projected, and the cell mask is created by manual thresholding. Using an opening operation with a disk-shaped structure (size = ~ 5 µm for the first step and ~ 2 µm for the second step), the protrusions are labelled. A minimal area condition is used to discard protrusions (~ 25 µm$^2$ for the first step and ~5 µm$^2$ for the second step) that are negligibly small. The final image shows the overlay of the protrusions (yellow for first step and orange for second step) and the cell body (light blue). Scalebar is 20 µm.*

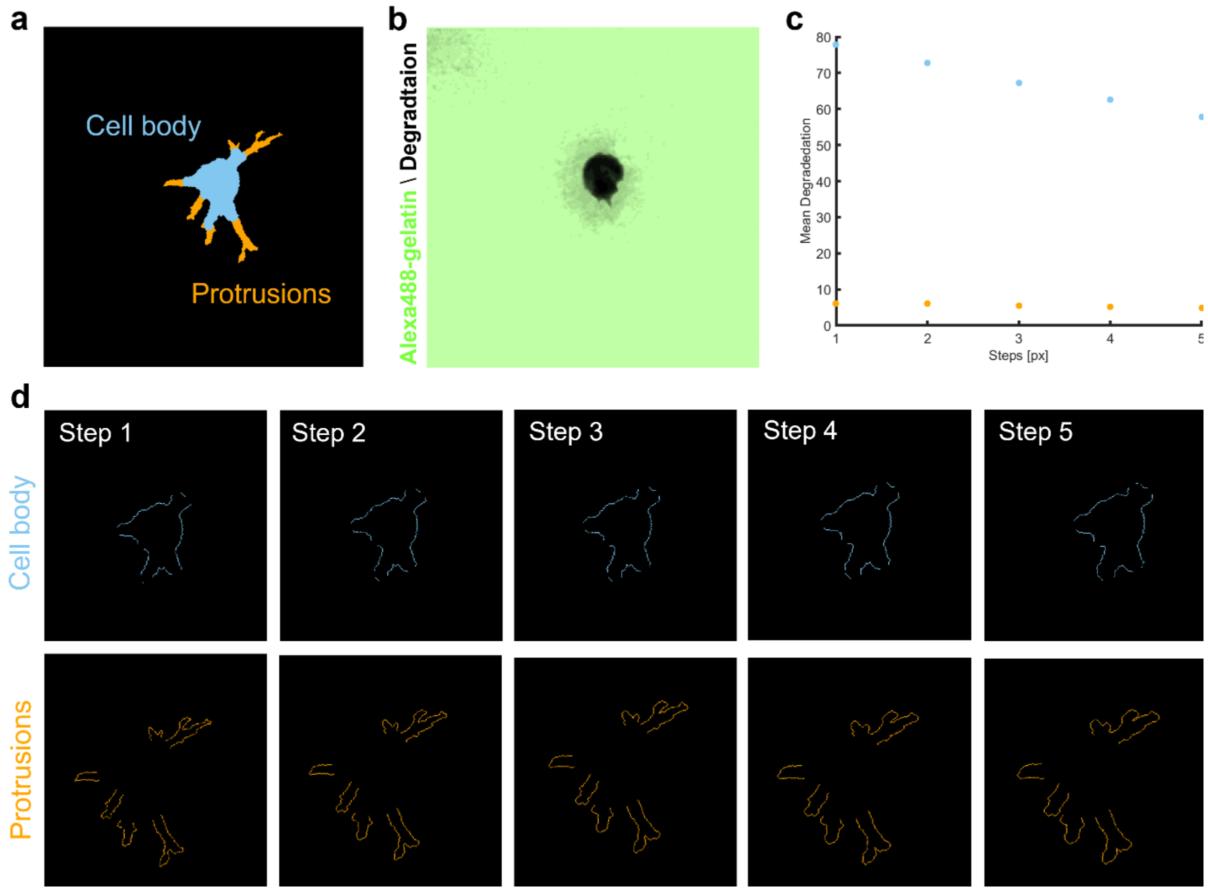


*Figure S7. Degradation quantification method for Fig. 3l. Degradation was assessed separately around the cell body and the protrusions within step segmentations. The step segmentations were generated by dilating the cell body / protrusion segmentation with a disk-shaped structure element with the size of 1 pixel. For each step, the mean degradation intensity within the step segmentation is reported. This is repeated for 30 steps (equivalent to a perimeter of 8µm).* ***a*** *Cell body and protrusion mask,* ***b*** *gelatin-488 minimum intensity projection,* ***c*** *Mean degradation curve for five steps,* ***d*** *Segmentation of 5 steps for the cell body and protrusions.*

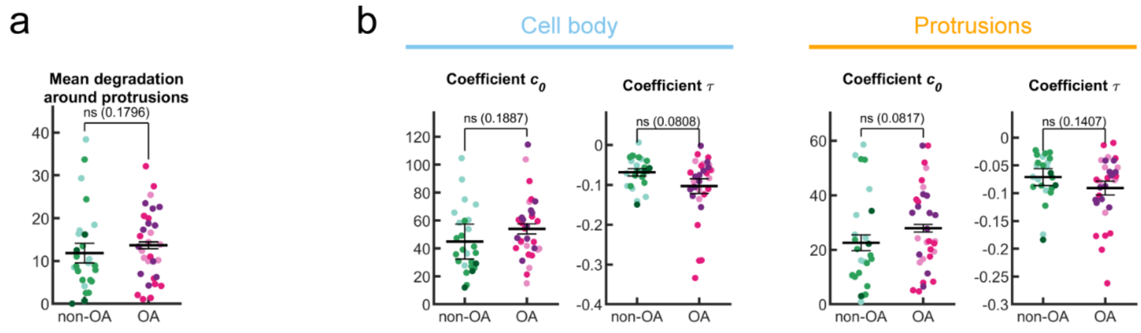


*Figure S8.* ***a*** *Overall mean degradation intensity around the protrusions within a perimeter of 5 µm: no difference was observed between non-OA and OA cells* ***b Analysis of the degradation curve decay profile:*** *coefficients retrieved from fitting an exponential curve $y = c_0 * e^{\tau * x}$ to the degradation intensity curve (Fig. 3l) of the cell body and protrusions. Coefficient $c_0$ represents the value at the cell boundary, coefficient $\tau$ represents the exponential decay constant, and x is the distance (in microns) from the cell boundary. No significant differences were observed between non-OA and OA cells regarding the coefficients for both the cell body and protrusions.*

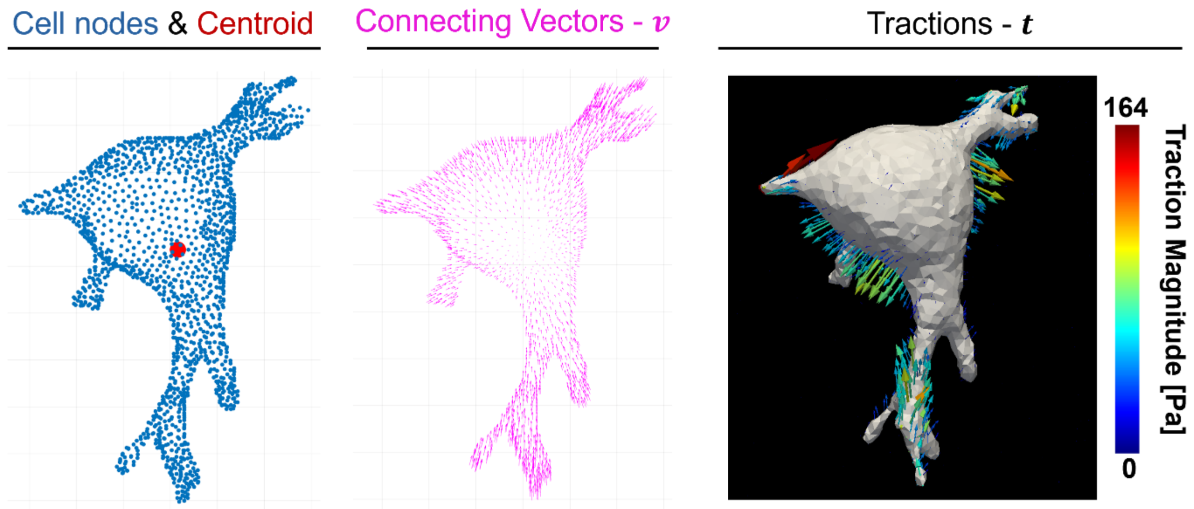


*Figure S9. Quantification of contractility, illustrated for a cell in 3D. Contractility is computed according to Eq. 1. A connecting unit vector pointing towards the mesh centroid is computed for each node on the cell surface mesh. Tractions in each node are projected on its corresponding connecting unit vector and contractility is computed by summing the (signed) magnitude of these projections for all nodes. The same projection is used to discriminate between pulling (projection pointing towards the centroid) and pushing tractions and displacements (projection pointing away from the centroid).*

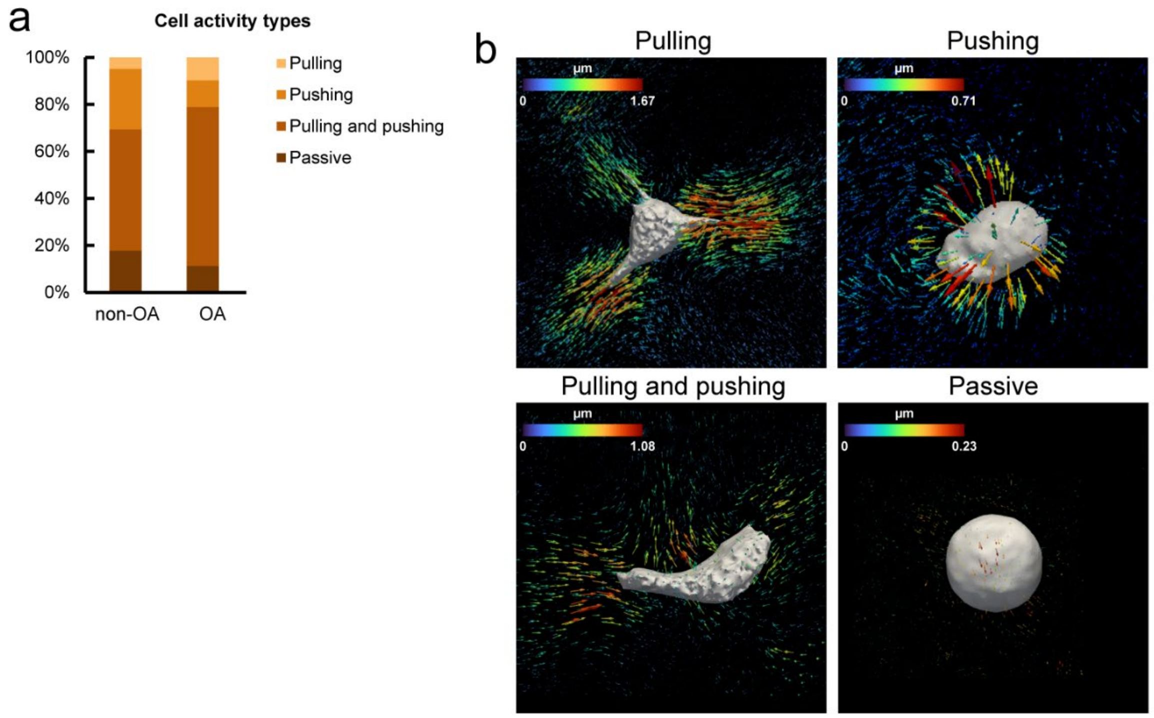


*Figure S10. **a** The proportion of cells categorized by their mechanical activity type: pulling, pushing, both pulling and pushing, or passive. **b** Representative images of different cell mechanical activity types, showing cell-generated matrix displacements.*
